# Tryptophan-substitution antimicrobial peptide temporin-1CEb: in vitro and in vivo antibacterial activity against clinically isolated multidrug-resistant *Klebsiella pneumoniae*

**DOI:** 10.64898/2025.12.30.696967

**Authors:** Fengquan Jiang, Yanjun Ma, Yunfei Zhang, Dejing Shang, Weibing Dong

## Abstract

Tryptophan (Trp)-substitution antimicrobial peptides (AMPs) exhibit enhanced interactions with bacterial cell membranes, potentially improving their antimicrobial efficacy. *K. pneumoniae* (20.59% of 2054 ICU isolates) is resistant to multiple clinically used antibiotics and presents significant treatment challenges. In the present study, three Trp-modified peptides (I4W, L12W, and I1WL5W) were generated by substituting Ile or Leu residues in temporin-1CEb, a peptide derived from frog skin, with Trp at various sites to assess their antibacterial effects and mechanisms against *K. pneumoniae*. Compared with L12W, both I4W and I1WL5W displayed superior antimicrobial activity and lower cytotoxicity. Mechanistic studies revealed that the AMPs exerted antibacterial and bactericidal effects through bacterial surface charge neutralization, insertion into bacterial cell membranes, increased permeability of both the inner and outer membranes, and disruption of membrane integrity. Notably, I1WL5W exhibited the most potent membrane-disrupting activity. Assessment of the impact of Trp-containing peptides on bacterial biofilms revealed that these peptides not only inhibited exopolysaccharide production and biofilm formation but also degraded preformed biofilms. A murine lung infection model was established to investigate the therapeutic efficacy of I1WL5W against MDRKP 1203-induced lung infection in mice. Compared with the control treatment, treatment with I1WL5W resulted in reduced bacterial counts and levels of IL-6 and TNF-α in both the blood and lung tissues of MDRKP 1203-infected mice, and treatment with I1WL5W improved lung tissue structure. The present study provides valuable insights for designing Trp-containing peptides with potent antimicrobial properties by facilitating their penetration across bacterial membranes.

## 1. Introduction

The emergence of antimicrobial resistance, driven by widespread antimicrobial utilization since the late 20th century, has precipitated a global public health crisis through accelerated evolution of multidrug-resistant (MDR) bacterial pathogens[1]. *K. pneumonia*e, a ubiquitous gram-negative opportunistic pathogen, colonizes environmental reservoirs, cutaneous surfaces, mucosal interfaces, and gastrointestinal ecosystems. This organism represents a critical nosocomial pathogen associated with severe clinical manifestations, including urinary tract infections, bacteremia, intra-abdominal infections, and ventilator-associated pneumonia [2, 3]. The increasing prevalence of multidrug-resistant strains, coupled with a limited arsenal of effective antibiotics, has contributed to increased mortality associated with gram-negative bacterial infections, particularly those caused by *K. pneumoniae*. There is an increasing trend in the clinical identification of carbapenem-resistant *K. pneumoniae*, prompting the creation of new treatment options and the investigation of strategies involving multiple drugs[4, 5]. Antimicrobial peptides (AMPs), which are typically composed of 8-15 amino acid residues, represent an early component of innate immunity across diverse organisms and serve as regulators of the adaptive immune system in higher eukaryotes[6, 7]. AMPs possess a range of biological functions, including antibacterial, antiviral, and cancer-fighting attributes, along with the capacity to prevent biofilm development and regulate immune reactions. Additionally, compared with traditional antibiotics, AMPs generally exhibit distinct advantages including minimal molecular mass, excellent solubility, low cytotoxic effects, and heat resistance[8]. However, the clinical translation of AMPs is hindered by challenges related to stability, production cost, toxicity, bioavailability, distribution, and metabolic stability[9–11]. Recent research endeavors have focused on enhancing AMP efficacy through structural modifications and synergistic combinations with other compounds. Elucidating the specific mechanism of action for each peptide is crucial for optimizing its therapeutic potential[12]. Consequently, the development of novel AMPs with simplified sequences, emphasizing key core residues, has been pursued[13].

Gram-negative and gram-positive bacteria exhibit anionic surface constituents—lipopolysaccharides (LPS) and teichoic acids, respectively—combined with negatively charged phospholipid bilayers. The cationic nature of AMPs facilitates electrostatic interactions with these bacterial surfaces[14]. This interaction contributes to the multifunctional and broad-spectrum antimicrobial activity of AMPs, positioning them as promising candidates for novel antimicrobial drug development[15, 16]. While classical models attribute AMP bactericidal effects to membrane permeabilization and cytosol leakage [17], contemporary studies have identified nonlytic mechanisms, including intracellular translocation without membrane compromise[18]. Tryptophan(Trp) residues, which are prevalent in natural AMPs, mediate membrane interfacial interactions via indole side chain hydrogen bonding (dipole moment ∼21 D) [19]. Building on the temporin-1CEb scaffold [20–22], we engineered Trp-substituted analogs through systematic replacement of Ile/Leu residues at positions 1, 4, 5, 11, and 12, generating mono-substituted (I1W, I4W, L5W, L11W, and L12W) and disubstituted (I1WL5W and I4WL5W) variants. Structure-activity analysis revealed enhanced gram-negative/-positive potency in I1W/I4W analogs versus attenuated activity in L11W/L12W derivatives. Di-substituted peptides exhibit expanded antimicrobial spectra, with positional Trp placement-rather than stoichiometry-emerging as the critical determinant of bactericidal efficacy[21, 22].

The present study evaluated the antibacterial efficacy of tryptophan-engineered antimicrobial peptides (I4W, L12W and I1WL5W) against multidrug-resistant *K. pneumoniae* (MDRKP 1203) and investigated their potential modes of action, including their impact on the inner membrane, the outer membranes, biofilm formation, capsule polysaccharide effects, and therapeutic potential in mice with pulmonary MDRKP 1203 infection. These findings establish an empirical foundation for the development of novel Trp-based AMP antimicrobial therapies that target nosocomial MDR pathogens.

## 2. Materials and Methods

### 2.1 Peptide

Trp-modified peptides (I4W, L12W, and I1WL5W) were commercially synthesized by GL Biochemistry Ltd. (Shanghai, China) with an HPLC-verified purity of ≥95%. Molecular mass validation was performed via matrix-assisted laser desorption/ionization time-of-flight mass spectrometry (MALDI-TOF MS; Shimadzu Corporation, Kyoto, Japan). The complete amino acid sequences and comprehensive physicochemical characterization of the parent peptide (Temporin-1CEb) and the engineered peptides are presented in **Table 1**. Net charge was predicted at physiological pH (7.4) by summing the charges of ionizable amino acid residues. The mean hydrophobicity values of peptides used in this study were calculated using hydrophobicity scales[23][24]. Amphipathicity values of all peptides were calculated by hydrophobic moment[25], using the software package JEMBOSS version 1.5[26].Helical wheel projections for the parent peptide (Temporin-1CEb) and the engineered peptides were generated using HeliQuest and are presented in Figure S1. The projections of the modified peptides (I4W, I1WL5W, and L12W) display a well-defined, continuous hydrophilic face (composed predominantly of basic, blue-colored residues) and a distinct hydrophobic face (formed by yellow-colored residues).

**Table 1.**
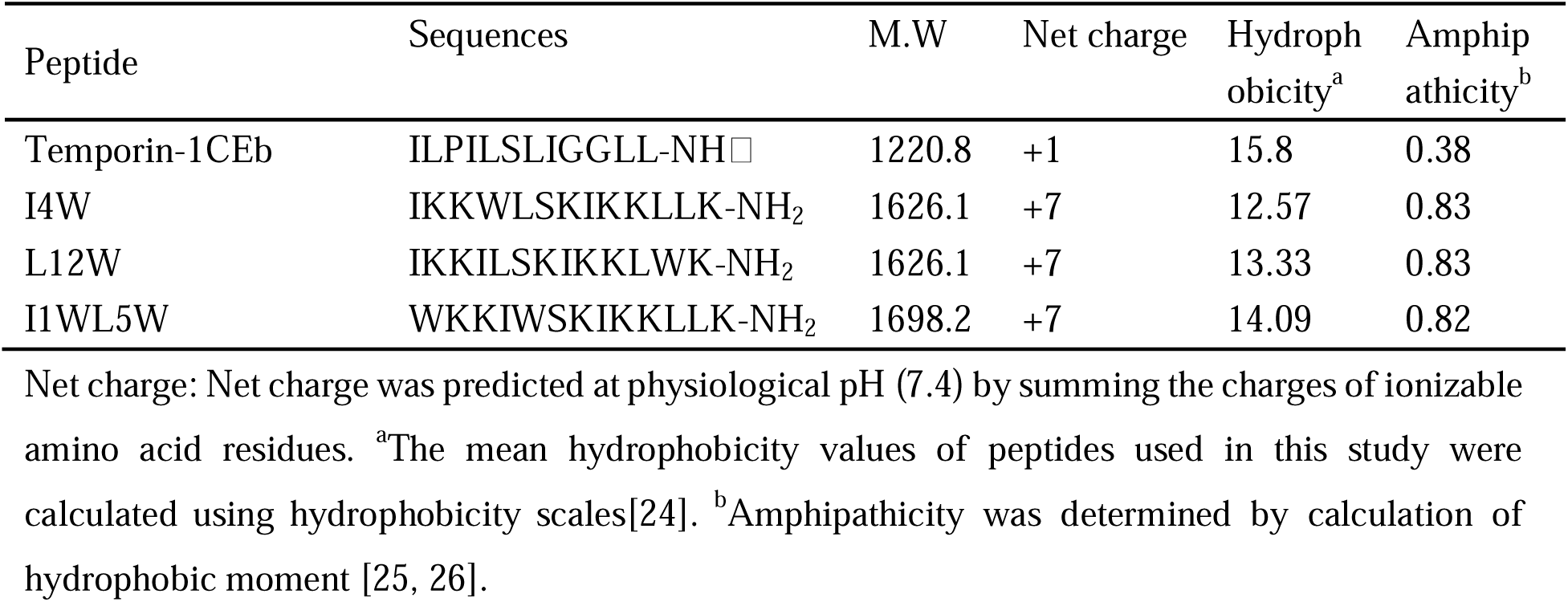
Amino-acid sequences and physicochemical properties.

### 2.2 Clinical microorganisms

Microorganisms was extracted from patients in the emergency ICU at Dalian Medical University’s First Affiliated Hospital in China, from January-December 2023. Specimens with suspected bacterial contamination and duplicates of the same strain cultured from the same patient and specimen type during the same time period were excluded, but specimens of the same strain cultured from the same patient but from different specimen types were included. Within a clinical laboratory setting, samples of sputum, urine, bronchial lavage, and secretions were added to various type of media, including blood agar, chocolate agar, and MacConkey agar plates. The inoculated media were subsequently incubated at a temperature range of 35-37°C for 18-24 hours. Blood culture specimens, collected during episodes of fever or chills, were dispensed into blood culture bottles containing anticoagulants (e.g., sodium citrate and heparin) and specialized growth media. The bottles were then incubated in an automated blood culture system. Upon detection of bacterial growth, immediate smear staining and subculturing onto appropriate solid media were performed[27]. In total, 49 strains of multidrug-resistant *K. pneumoniae*(MDRKP) were isolated and investigated. The MDRKP variants demonstrated resistance to some treatments, such as erythromycin, penicillin, piperacillin, azithromycin, and vancomycin.

### 2.3 Antibiotic susceptibility tests

Bacterial identification and antimicrobial susceptibility testing were conducted using the MicroScan WalkAway-96 Plus system (Siemens Healthcare, Germany) at the First Affiliated Hospital of Dalian Medical University, following the National Guide to Clinical Laboratory Procedures. The results were interpreted per the CLSI guidelines[28, 29]. Quality control utilized the *Staphylococcus aureus* ATCC 29213, *Escherichia coli* ATCC 25922, *Pseudomonas aeruginosa* ATCC 27853, and *Enterococcus faecalis* ATCC 29212 reference strains, which were provided by the Ministry of Health Clinical Laboratory Center. Antimicrobial susceptibility patterns were analyzed using WHONET 2020 software.

### 2.4 Determination of Minimal Inhibitory Concentrations (MICs)

Minimum inhibitory concentrations (MICs) were determined via standardized broth microdilution assays according to CLSI guidelines [28, 29]. The peptides were subjected to two-fold serial dilutions (200–1.56 μM) in sterile 96-well plates, with each well receiving 100 μL of peptide solution and 100 μL of bacterial inoculum (mid-log phase cultures adjusted to 2 × 10□ CFU/mL in Mueller-Hinton broth). The blank control was sterile Mueller-Hinton broth only and the positive control wells contained 100 μL of Mueller-Hinton broth without the peptide solution, along with 100 μL of the bacterial inoculum, to verify normal bacterial growth under the experimental conditions. Following 15 to 20 hours of incubation at 37°C. MIC values were defined as the lowest concentration of an antibiotic that blocks the growth of bacteria, eliminating any small haze or turbidity. Each experiment was carried out separately and three times to ensure accuracy.

### 2.5 Bactericidal kinetics

The bactericidal activity of the peptides toward the MDRKP 1203 strain was assessed via time-kill curves[30]. Logarithmic-phase cultures (10□ CFU/mL) were exposed to peptide concentrations (1×, 4×, 8×, and 16× MIC) under aerated incubation (37°C, 180 min). Following dual washing with sterile Mueller-Hinton broth and centrifugal pelleting (4,000 rpm, 10 min), processed samples underwent serial logarithmic dilutions (10²–10□) for quantitative culturing via automated spiral plating (Don Whitley Scientific, UK). The experimental controls included untreated bacterial suspensions and gentamicin (3.13 μM, 1×MIC). Viable colonies were enumerated post incubation to determine their bactericidal kinetics.

### 2.6 Determination of post-antibiotic effects (PAEs)

The PAE refers to the duration during which an antimicrobial drug continues to suppress microorganism growth after it has been removed from culture, even when the serum concentration of the antimicrobial has decreased below the minimum inhibitory concentration or has disappeared[30]. Postantibiotic effects (PAEs) were evaluated by exposing logarithmic-phase MDRKP 1203 cultures (10□ CFU/mL) to peptide concentrations of 1×, 2×, or 4× MIC during for 60 minutes at 37°C. After treatment, the bacterial suspensions were washed twice with Mueller-Hinton (MH) broth, pelleted via centrifugation (4,000 rpm, 10 minutes), resuspended in fresh medium, and incubated aerobically (37°C, 8 hours) with agitation. Serial dilutions of processed cultures were quantified using a spiral-plating system (Don Whitley Scientific, UK). Untreated bacterial cultures and gentamicin-treated samples served as negative and positive controls, respectively. PAE duration was calculated using the following equation: PAE duration = T_test_−T_control_; where T represents the time required for bacterial counts to increase by 1.0 log□□ CFU/mL posttreatment [30].

### 2.7 Scanning Electron Microscopy

Log-phase bacterial suspensions (1 × 10□ CFU/mL) were exposed to peptide concentrations of 1× or 2× MIC for 60 minutes at 37°C. Following three PBS washes, the bacterial pellets were subjected to primary fixation in 2.5% glutaraldehyde/0.1 M cacodylate buffer (4°C, 2 hours) and secondary fixation in 1% osmium tetroxide (2 hours). Sequential ethanol dehydration (graded concentrations) preceded lyophilization and gold sputter coating. Cellular ultrastructural alterations were analyzed using field-emission scanning electron microscopy (Hitachi SU8010, Japan) [31]. Untreated bacterial cultures served as negative controls.

### 2.8 Trp fluorescence and KI quenching assays

The tryptophan fluorescence emission spectra of peptide-treated MDRKP 1203 cultures were analyzed using established quenching methodologies [32]. The bacterial cells that were grown to logarithmic-phase were pelleted by centrifugation (4,000 ×g for 10 min), washed twice with sterile phosphate-buffered saline (PBS, pH 7.4) to remove any residual culture medium and unbound peptides, and finally resuspended in an equal volume of clear PBS for fluorescence measurement. Subsequently, logarithmic-phase bacteria (1×10□ CFU/mL) were incubated with various peptide concentrations in 96-well plates (25°C, 10 min–3 h). The fluorescence of the peptide was subsequently determined at excitation/emission wavelengths of 280/300-400 nm using a Varioskan Flash microplate reader (Thermo Scientific, Beijing, China).

Collisional quenching experiments employed potassium iodide (KI) as an aqueous-phase quencher. Peptide-treated MDRKP 1203 cultures (1×10^5^ CFU/mL, 1-hour incubation) were exposed to a gradient range of KI concentrations (0, 20, 40, 60, 80, 100, 120, and 140 mM). The fluorescence of the peptide was determined at excitation/emission wavelengths of 295/350 nm to eliminate interference from the absorbance of the quencher[33]. The quenching efficiency was calculated via the Stern-Volmer equation as follows: F_0_/F= 1+*K_SV_*[Q]; where F_0_ and F are the fluorescence intensities in the presence and absence of the quencher, respectively; *Ksv* denotes the Stern-Volmer constant; and [Q] indicates the quencher molar concentration.

### 2.9 Inner Membrane Depolarization Assay

Bacterial membrane depolarization was quantified using the potential-sensitive fluorescent probe DiSC3-5 [33]. The logarithmic-phase cultures were pelleted (3,000 ×g, 10 minutes), washed twice in HEPES buffer (5 mM, pH 7.2) containing 20 mM glucose and 100 mM KCl, and adjusted to 2 × 10□ CFU/mL in assay buffer. Bacterial suspensions were aliquoted into black-walled microplates containing 4 μM DiSC3-5, with baseline fluorescence (λ_ex_=622 nm/λ_em_=670 nm) monitored at 30-second intervals using a Varioskan Flash microplate reader (Thermo Fisher Scientific, China). Fluorescence stabilization confirmed maximal dye incorporation into polarized membranes. Test peptides (1×, 2×, and 4× MIC) were introduced, and fluorescence changes were recorded relative to those of the controls (0.1% Triton X-100) for complete depolarization and vancomycin as a nondepolarizing negative control.

### 2.10 Outer Membrane Permeability Assay

Outer membrane permeability was assessed using 1-N-phenylnaphthylamine (NPN) uptake assays following established protocols[34]. The logarithmic-phase bacterial cultures were pelleted via centrifugation (3000 ×g 10 minutes), washed two times with 5 mM HEPES buffer, and resuspended in 10 mM sodium phosphate buffer (pH 7.4). Bacterial suspensions (2 × 10□ CFU/mL) were incubated with 10 μM NPN in the presence of peptide gradients (1.56–100 μM) for 1 hour at 25°C. Fluorescence emission (λ_ex_=350 nm/λ_em_=420 nm) was quantified using a Varioskan Flash microplate reader (Thermo Fisher Scientific). The experimental controls included vancomycin (negative) and gentamicin (positive). All measurements were performed in triplicate for each peptide concentration.

### 2.11 Liposome preparation

Lipid components (phosphatidylglycerol [PG], phosphatidylcholine [PC], phosphatidylethanolamine [PE], cardiolipin [CL], cholesterol) were commercially sourced from Sigma-Aldrich (China). Lipopolysaccharide (LPS) was extracted from MDRKP 1203 cultures via hot phenol-water purification. Synthetic liposomes were formulated to replicate gram-negative bacterial membrane architectures as follows: inner membrane (PE/PG/PC at a 0.15:1.85:1 mass ratio)[35] and outer membrane (PC/PG/PE/LPS at a 5:2.95:1.05:1 mass ratio)[36]. Calcein-encapsulated liposomes were synthesized through freeze-thaw-sonication[22]. Lipid mixtures were dissolved in chloroform/methanol (2:1 v/v), followed by organic solvent evaporation under reduced pressure. Hydration with 90 mM calcein solution (20 mM TES and 100 mM NaCl, pH 7.4) preceded 10 freeze-thaw cycles (−196°C/50°C). Liposomal suspensions were extruded through 200-nm polycarbonate membranes (10 cycles) and purified via size-exclusion chromatography (Sephadex G-50 column).

### 2.12 Leakage of calcein from liposomes

Calcein-loaded liposomes were exposed to peptide concentrations (1.56–50μM) for 30 minutes, and fluorescence emission (λ_ex_=485 nm/λ_em_=530 nm) was quantified via a Varioskan Flash microplate reader to assess membrane permeabilization. The percentage of calcein release was calculated using the following formula: calcein release (%) = (F-F_0_)/(F_100_-F_0_) × 100%; where F and F_0_ represent the fluorescence intensity of the calcein-labeled liposomes in the presence or absence of peptides, respectively; And F_100_ is the fluorescence intensity of the calcein-labeled liposomes treated with 10%(w/v) Triton X-100[37]. Vancomycin and gentamicin served as negative and positive controls, respectively.

### 2.13 Confocal laser scanning microscopy

Membrane integrity was measured using the LIVE/DEAD^®^ BacLight™ Bacterial Viability Kit (Invitrogen, China) according to the manufacturer’s protocols. Logarithmic-phase MDRKP 1203 cultures were exposed to 12.5 μM or 25 μM peptides (37°C, 1 h) and dual-stained with SYTO9 and propidium iodide (PI) under dark conditions (30 minutes), Images were then acquired via confocal laser scanning microscopy (LSM 710, Zeiss, Germany). Viable cells exhibited SYTO9-associated green fluorescence (λ_ex/em_=480/500 nm), whereas membrane-compromised cells displayed PI-induced red fluorescence (λ_ex/em_=490/635 nm)[30].

### 2.14 Zeta potential measurements

Zeta potential measurements were conducted on mid-logarithmic-phase MDRKP 1203 cultures (2 × 10□ CFU/mL) using disposable zeta potential cuvettes with gold-coated electrodes at 25°C. The bacterial suspensions were titrated with increasing peptide concentrations (1.56–100 μM) and equilibrated for 30 minutes prior to analysis. Surface charge quantification was performed using a Zetasizer Nano ZS (Malvern Panalytical, UK) equipped with a 633-nm HeNe laser. Five technical replicates (100 runs each) were analyzed per concentration. The final zeta potential values represent the means of 15 independent measurements per experimental condition. Gentamicin served as the positive control for membrane charge modulation [38].

### 2.15 Biofilm Formation Assay

MDRKP 1203 bacterial cultures in mid-logarithmic growth phase (1 × 10□ CFU/ml) were added into the wells of a 96-well microplate and incubated with peptide solutions prepared at serial dilutions corresponding to 1/8, 1/4, 1/2, 1×, and 2× the minimum inhibitory concentration (MIC) for 18 hours at 37°C under static conditions. Following incubation, the medium containing non-adherent cells were meticulously aspirated. The remaining adherent biofilms were then fixed with methanol for 15 minutes, followed by staining with a 0.1% (mass/volume) aqueous crystal violet solution for 5 minutes. After staining, the plates were subjected to three sequential washes with deionized water to eliminate unbound dye. Biofilm-associated crystal violet was solubilized via immersion in ethanol (95% v/v) for 30 minutes. Optical density quantification at 590 nm was performed using a Varioskan Flash spectral scanning multimode reader (Thermo Fisher Scientific, Beijing). The experimental controls included untreated LB broth (negative control) and gentamicin-treated samples (positive control). The biofilm inhibition efficacy was calculated through the following equation:[(OD590_peptide_ - OD590_control_)/OD590_control_] × 100%. The MBIC_50_, or the minimum biofilm inhibitory concentration, was identified as the lowest peptide concentration capable of preventing biofilm formation by 50% or more[39, 40]. Shang et al. outlined a method for assessing peptide effectiveness against biofilms aged one day[41]. Biofilms from the MDRKP 1203 strain, which were aged one day, were washed with PBS and exposed to varying peptide concentrations at 37°C for one day. The biofilms were subsequently fixed, stained, and measured as previously described. MBRC50, or the minimum biofilm reduction concentration, was identified as the minimal peptide level required to decrease the biofilm by 50% or more. GM served as the positive control.

### 2.16 Determination of the Exopolysaccharides in Biofilms

Exopolysaccharide quantification was performed via the phenol-sulfuric acid method [42]. Mid-log phase MDRKP 1203 cultures (1 × 10^5^ CFU/mL) were treated with subinhibitory-to-inhibitory peptide concentrations (1/4, 1/2, and 1× MIC) under static incubation (37°C, 24 hours). Bacterial pellets were harvested by centrifugation (10,000 ×g, 15 minutes), resuspended in high-ionic-strength buffer (10 mM KPO_4_, 5 mM NaCl, and 2.5 mM MgSO_4_, pH 7.0), and recentrifuged (10,000 ×g, 30 minutes). The supernatants were mixed with three volumes of absolute ethanol for exopolysaccharide precipitation (4°C, 16 hours). The precipitated polysaccharides were reacted with 5% phenol and concentrated sulfuric acid, and the absorbance was quantified at 490 nm. Untreated cultures (LB broth) and gentamicin-treated samples served as negative and positive controls, respectively.

### 2.17 Establishment of the MDRKP 1203-induced pulmonary infection model

Six-week-old male KM mice (22-24 g, sourced from the Laboratory Animal Center of Dalian Medical University) were utilized in live studies[31]. The mice received a daily intraperitoneal injection of cyclophosphamide (100 mg/kg body weight) over a period of 3 days to trigger immunosuppression before infection. The mice were then anesthetized through an intraperitoneal injection of 10% chloral hydrate (40 mg/kg body weight). The nasal drip method was used to induce the pulmonary infection model. A mid-log-phase MDRKP 1203 suspension (1 × 10^7^ CFU/mL in 30 μL total volume) was administered intranasally using a calibrated micropipette. The suspension was delivered slowly in 5 μL aliquots alternating between nostrils, with 5 second intervals between drops to allow for complete inhalation during spontaneous breathing to draw the suspension into the lower respiratory tract and lungs. The mice were randomly divided into the following six groups (n=12/group): the untreated group (control group); the infected model group with 0.9% NaCl; the I1WL5W group (2, 4, and 8 mg/kg body weight, respectively); and the gentamicin group (Gen, 2 mg/kg body weight). Twenty-four hours after bacterial injection, various doses of I1WL5W, saline, or gentamicin were administered intraperitoneally (i.p.). The drugs were administered daily to the mice over a span of 9 days. The vital signs and body weights were recorded daily. Following the administration of peptides, on Days 3, 6, and 9, the mice were euthanized, and their blood and lung tissues were harvested to assess bacterial colonization and release of inflammatory factors. These tissues were then stained with hematoxylin and eosin (H&E) for histological examination. All procedures involving animals were approved by the Animal Use and Care Committee of Liaoning Normal University (ethical approval number: LL2023063).

### 2.18 Bacterial counts in mouse blood and lung

The bacterial counts in the blood and lungs of mice were determined using a spiral-plating system [31]. In brief, the lung tissue was homogenized in 1 mL of PBS, and the homogenates were diluted, plated on MH agar and incubated overnight at 37°C. The bacterial colonies present in both the lungs and blood samples were counted.

### 2.19 Inflammatory factor assay

Using enzyme-linked immunosorbent assay (ELISA) kits (RapidBio, Shanghai, China), the concentrations of the tumor necrosis factor α (TNF-α) and interleukin-6 (IL-6) proinflammatory agents were measured in blood and lung tissues according to the manufacturer’s guidelines [31]. In brief, lung tissues were homogenized in 1 mL of PBS and centrifuged at 10,000 rpm for 15 minutes. The TNF-α and IL-6 concentrations in the supernatant were then assessed.

### 2.20 Histological analysis

Lung samples were fixed in 4% formaldehyde, embedded in paraffin, sectioned, and stained with hematoxylin and eosin (H&E)[31]. The morphological alterations in the lungs were examined using a light microscopy (40×, Olympus, Japan).

### 2.21 Statistical Analysis

Each experiment was replicated three times. The outcomes are presented as averages and standard errors. Significance was assessed using the paired Student’s t-test or one-way ANOVA. p < 0.05 was defined as a statistically significant.

## 3. Results

### 3.1 Multidrug-resistant *K. pneumoniae* in the emergency ICU (eICU)

In total, 2,176 strains of pathogenic microorganisms were isolated and cultured from various types of specimens collected in the eICU of the First Affiliated Hospital, Dalian Medical University from January 2023 to December 2023. After excluding duplicates of the same strain cultured from the same patient and specimen type, 2054 strains met the inclusion criteria. The specimen types with the three highest detection rates were sputum (75.7%), blood (7.6%), and bronchial lavage fluid (6.8%) (Table **S1**). The pathogenic microorganisms detected in the eICU were mainly gram-negative bacteria(65.0%), followed by fungi (26.1%) and gram-positive bacteria (8.9%). Among the 182 strains of gram-positive bacteria, *Staphylococcus aureus* accounted for the greatest proportion (2.78%), followed by *Enterococcus faecalis* (1.9%), *Staphylococcus hominis* (1.36%) and *Staphylococcus epidermidis* (1.12%). Among the 1335 gram-negative bacteria, *K. pneumoniae* accounted for the greatest proportion (20.59%), followed by *Acinetobacter baumannii* (19.43%)*, Stenotrophomonas maltophilia* (6.86%) and *Pseudomonas aeruginosa* (6.23%)(**Table S2-3**).

*K. pneumoniae* isolates exhibited elevated resistance to first-/second-generation cephalosporins (70–100%), with the exception of cefotetan (26.38%), and they demonstrated limited resistance to amikacin (11.3%), polymyxins (<4%), and tigecycline (<4%). The carbapenem resistance rates (imipenem/meropenem) were approximately 70%. *Escherichia coli* displayed reduced carbapenem resistance (<17%) alongside low resistance to amikacin, polymyxins, and cefotetan. *Acinetobacter baumannii* demonstrated near-universal carbapenem resistance (>95%) but retained susceptibility to tigecycline/polymyxins (<2%). *Pseudomonas aeruginosa* exhibited moderate carbapenem resistance (≈40%) and preserved susceptibility to amikacin/cefepime (<8%). The fungal isolates were predominantly *Candida albicans*, demonstrating full susceptibility to amphotericin B/5-fluorocytosine and limited resistance to azoles (fluconazole/itraconazole <25%).

### 3.2 Antibacterial activity of Trp-containing peptides against multidrug resistant *K. pneumoniae*

The minimum inhibitory concentrations (MICs) of tryptophan-containing peptides (I4W, L12W, and I1WL5W) were determined against 49 clinical multidrug-resistant *Klebsiella pneumoniae* (MDRKP) isolates from the First Affiliated Hospital of Dalian Medical University. As shown in **Table 2**, I1WL5W exhibited antibacterial activity against 27 multidrug-resistant *K. pneumoniae* strains, with an MIC of 50 μM accounting for 55.1% of all strains. In addition, L12W exhibited antibacterial activity against 22 strains, with an MIC of more than 200 μM accounting for the greatest percentage (44.9%), and I4W exhibited antibacterial activity against 14 strains, with MICs of 100 and 200 μM accounting for the greatest percentage (28.57%). The overall inhibitory effect of I1WL5W on 49 clinically isolated *K. pneumoniae* strains was better than that of L12W and I4W. All three Trp-containing peptides, namely, I4W, L12W and I1WL5W, showed high antibacterial activity against the multidrug-resistant *K. pneumoniae* 1203 (MDRKP 1203) strain, with MICs of 12.5, 25 and 12.5 µM, respectively. Based on these findings, the MDRKP 1203 strain was selected as the target pathogen for subsequent experiments.

**Table 2.** MICs of the peptides against 49 strains of *K. pneumoniae* isolated from clinical practice.

| *K. pneumoniae*<br>strain | MIC (μM) |  |  |
| --- | --- | --- | --- |
|  | L12W | I4W | I1WL5W |
| 1 | 100 | 100 | 50 |
| 2 | > 200 | > 200 | 50 |
| 3 | > 200 | > 200 | 100 |
| 4 | 25 | > 200 | 50 |
| 5 | > 200 | 200 | 50 |
| 6 | 25 | 12.5 | 50 |
| 7 | > 200 | 200 | 12.5 |
| 8 | > 200 | > 200 | 50 |
| 9 | > 200 | 200 | 50 |
| 10 | > 200 | 200 | 50 |
| 11 | > 200 | > 200 | 50 |
| 12 | > 200 | > 200 | 50 |
| 13 | > 200 | 100 | 50 |
| 14 | 100 | 100 | 50 |
| 15 | 25 | 50 | 12.5 |
| 16 | > 200 | 200 | 50 |
| 17 | > 200 | 200 | 100 |
| 18 | 200 | 100 | 50 |
| 19 | 50 | 12.5 | 100 |
| 20 | > 200 | 200 | 100 |
| 21 | 200 | 200 | > 200 |
| 22 | 200 | 200 | > 200 |
| 23 | 50 | 200 | 100 |
| 24 | 50 | 50 | 50 |
| 25 | 100 | 25 | 6.25 |
| 26 | > 200 | 200 | 50 |
| 27 | 25 | 100 | 12.5 |
| 28 | 200 | 100 | 50 |
| 29 | > 200 | 200 | 50 |
| 30 | 200 | 200 | 100 |
| 31 | > 200 | > 200 | 100 |
| 32 | 200 | 100 | 50 |
| 33 | 200 | > 200 | 50 |
| 34 | > 200 | > 200 | 100 |
| 35 | 200 | 100 | 50 |
| 36 | 200 | 100 | 50 |
| 37 | 25 | 50 | 25 |
| 38 | > 200 | 200 | 50 |
| 39 | > 200 | > 200 | 50 |
| 40 | > 200 | > 200 | 50 |
| 41 | 100 | 100 | 25 |
| 42 | 100 | 100 | 25 |
| 43 | 100 | 50 | 50 |
| 44 | > 200 | 100 | 25 |
| 45 | > 200 | > 200 | 100 |
| 46 | 200 | > 200 | 12.5 |
| 47 | 100 | 100 | 25 |
| 48 | 200 | 100 | 50 |
| 49 (MDRKP1203) | 25 | 12.5 | 12.5 |

MIC: minimal peptide concentration required for total inhibition of microbe growth in liquid medium. The values are the means of three independent experiments performed in four replicates

### 3.3 Bactericidal kinetics and postantibiotic effects (PAEs) of Trp-containing peptides on the MDRKP 1203 strain

The bactericidal temporal dynamics of I4W, L12W, and I1WL5W demonstrated concentration-dependent microbial eradication profiles. At 16 ×MIC, all peptides achieved complete microbial eradication within 180 minutes, exhibiting greater than 4log10 viability suppression within the initial 60-minute exposure (**Figure 1A-C**). Comparative analysis revealed that the microbial inactivation dynamics of gentamicin at 1×MIC were comparable to those of peptide formulations at equivalent concentrations. Rapid bacteriolytic kinetics correlated with prolonged postantibiotic effects, manifesting sustained suppression durations of 4 h (I4W), 4 h (L12W), and 4 h (I1WL5W) after exposure for 60 minutes at 1×MIC (**Figure 1D-F**), which indicated irreversible membrane-targeted bactericidal mechanisms.

**Figure 1.**
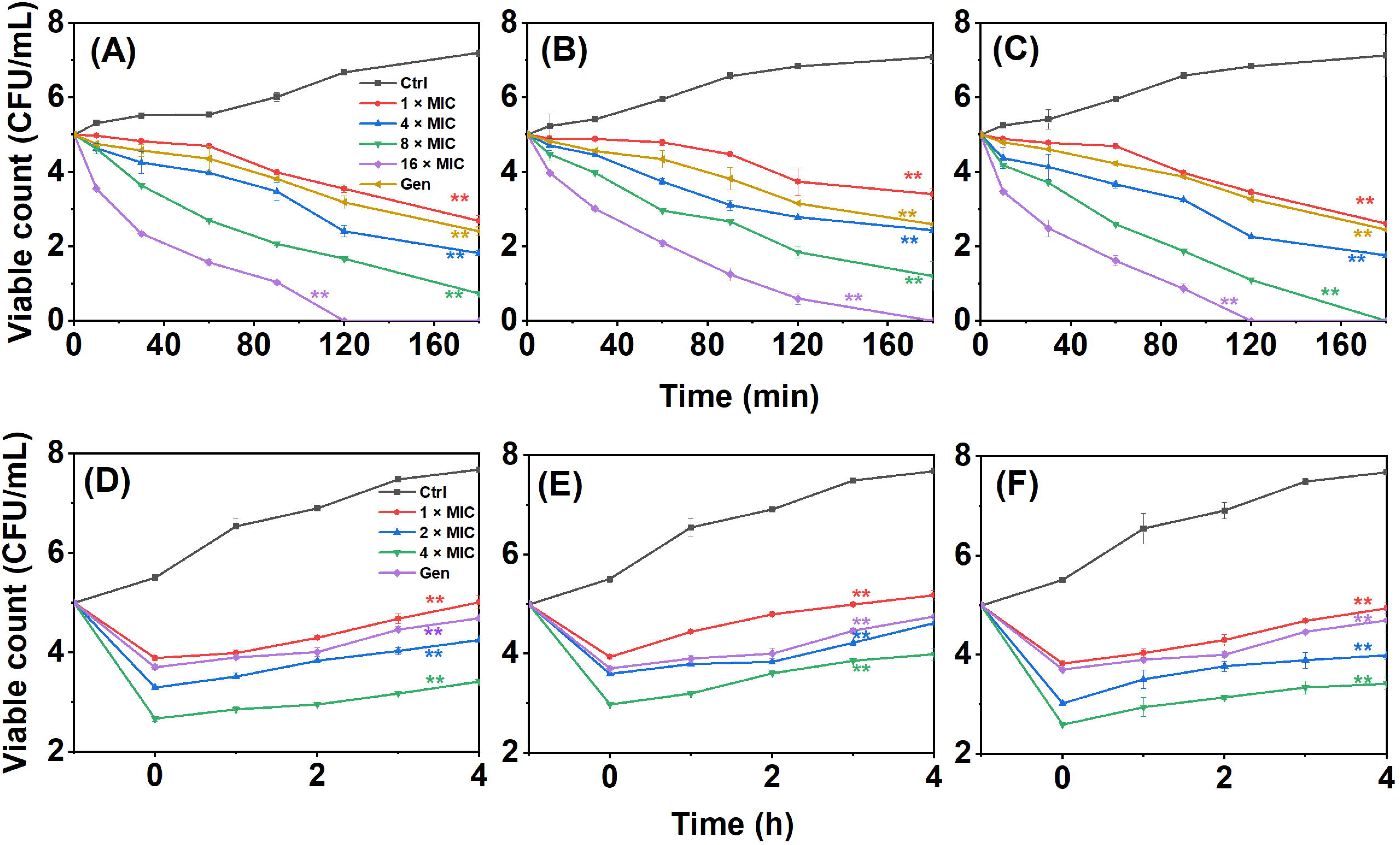
Time-killing curve for Trp-containing AMPs against MDRKP 1203 (**A** for I4W, **B** for L12W, **C** for I1WL5W). MDRKP 1203 cell suspensions were incubated with peptides at the final concentration of 1×, 4×, 8×, 16 × MIC for 1-3 h, n = 3. The surviving bacteria were counted after being diluted. Bacteria suspensions treated without the peptides were as negative control and Gen was as positive control. The PAEs of AMPs against MDRKP 1203 (**D** for I4W, **E** for L12W, **F** for I1WL5W), n = 3. Data were expressed as the mean ± SEM by two-way ANOVA; \*\**p* < 0.01 versus control.

### 3.4 Membrane damage activity of Trp-containing peptides in MDRKP1203 cells

Ultrastructural alterations in MDRKP 1203 pathogens following tryptophan-peptide exposure were characterized through scanning electron microscopy (**Figure 2A**). Pathogen cohorts subjected to different peptide concentrations exhibited profound morphological derangements, manifesting pronounced surface texturing with concave deformations accompanied by compromised envelope integrity. These structural disintegrations progressed to complete cellular lysis with concomitant cytoplasmic constituent extrusion, demonstrating concentration-dependent membrane destabilization efficacy. These results indicated that the peptides disrupted the bacterial membranes and led to a loss of barrier function.

**Figure 2.**
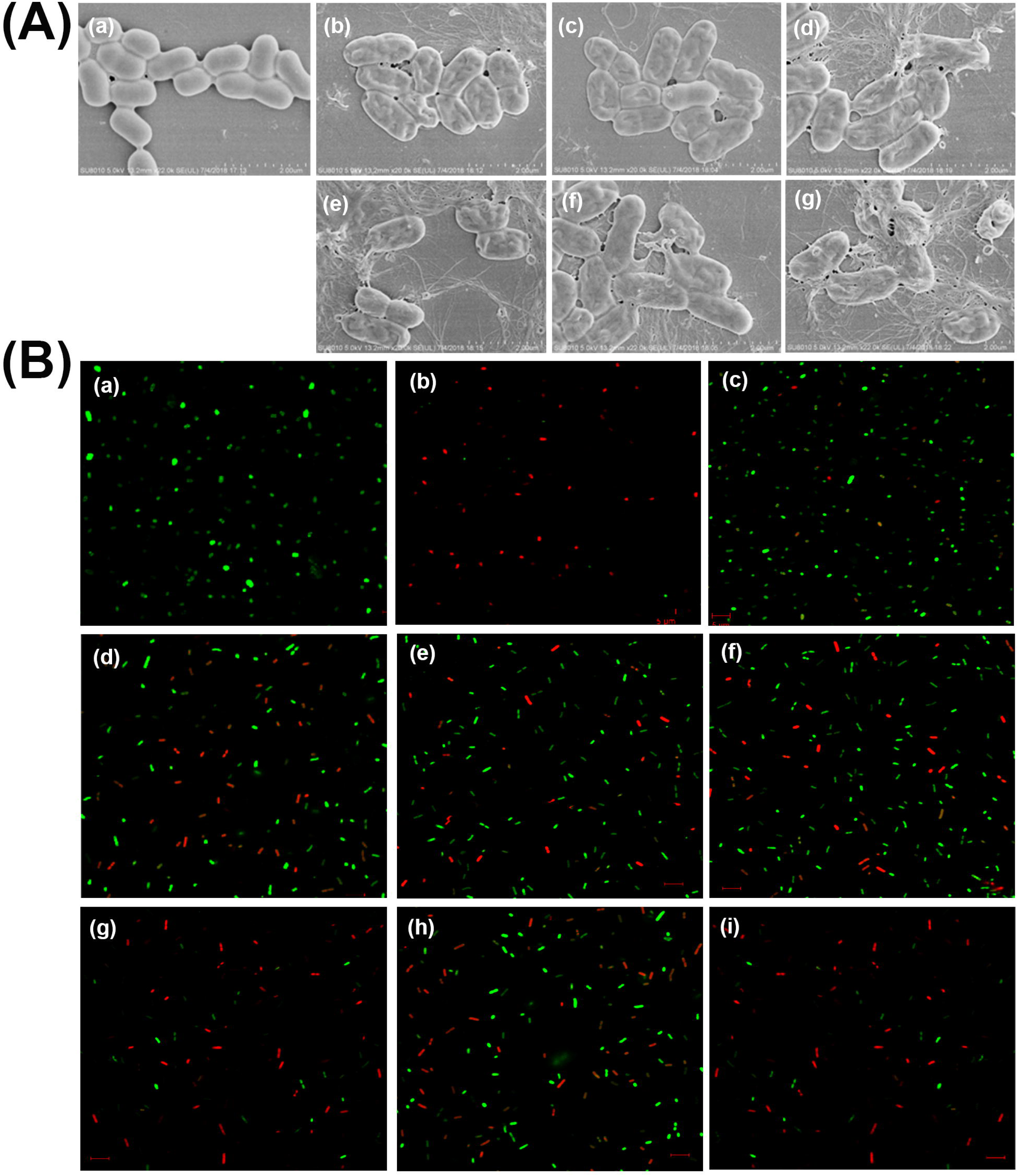
(**A**) Scanning electron micrographs of MDRKP 1203 without and with peptide treatment. a: Untreated control; b-d: Treated with 1×MIC of I4W, L12W, and I1WL5W, respectively; e-g: Treated with 2×MIC of I4W, L12W, and I1WL5W, respectively. (**B**) Live/dead staining images of MDRKP 1203 exposed to I4W, L12W, and I1WL5W. a: Untreated control; b-f: Treated with 12.5 μM of gentamicin, vancomycin, I4W, L12W, and I1WL5W, respectively; g-i: Treated with 25 μM of I4W, L12W, and I1WL5W, respectively.

A membrane integrity assessment employing dual fluorochrome labeling was conducted via confocal microscopy. SYTO 9, a membrane-permeant fluorophore, selectively labels viable cells with green emission through unimpaired membrane integration, whereas propidium iodide (PI) exclusively infiltrates structurally compromised membranes, resulting in red fluorescence (**Figure 2B**). Untreated MDRKP 1203 populations maintained green chromatic signatures, whereas peptide-exposed samples (12.5-25μM, 60 min) demonstrated complete chromatic transition to red emission spectra, confirming that membrane permeabilization was sufficient for PI internalization. Comparative analysis revealed that I4W and I1WL5W induced more pronounced membrane destabilization than L12W at a concentration of 25μM. The quantitative fluorometric ratios indicated that peptide-treated samples presented intermediate SYTO9/PI signal intensities between those of vancomycin-treated (lower disruption) and gentamicin-exposed (higher disruption) counterparts at equivalent dosage levels.

Fluorescence spectral profiling of tryptophan-enriched peptides revealed that the membrane interaction dynamics were mediated by indole-mediated hydrophobic anchoring. Trp residue emission spectra (λmax ≈340 nm in the aqueous phase) revealed changes in the bathochromic emission profile upon complexation with MDRKP 1203 cells (**Figure 3A**). In contrast, progressive microbial titration induced hypsochromic spectral displacement, confirming membrane localization through fluorophore environmental shielding. Quantitative analysis revealed that I4W, L12W, and I1WL5W presented maximal wavelength shifts of 26 nm, 28 nm, and 32 nm respectively during bacterial complexation. These hypsochromic transitions were directly correlated with the depth of indole group insertion into the hydrophobic domains of the membrane.

**Figure 3.**
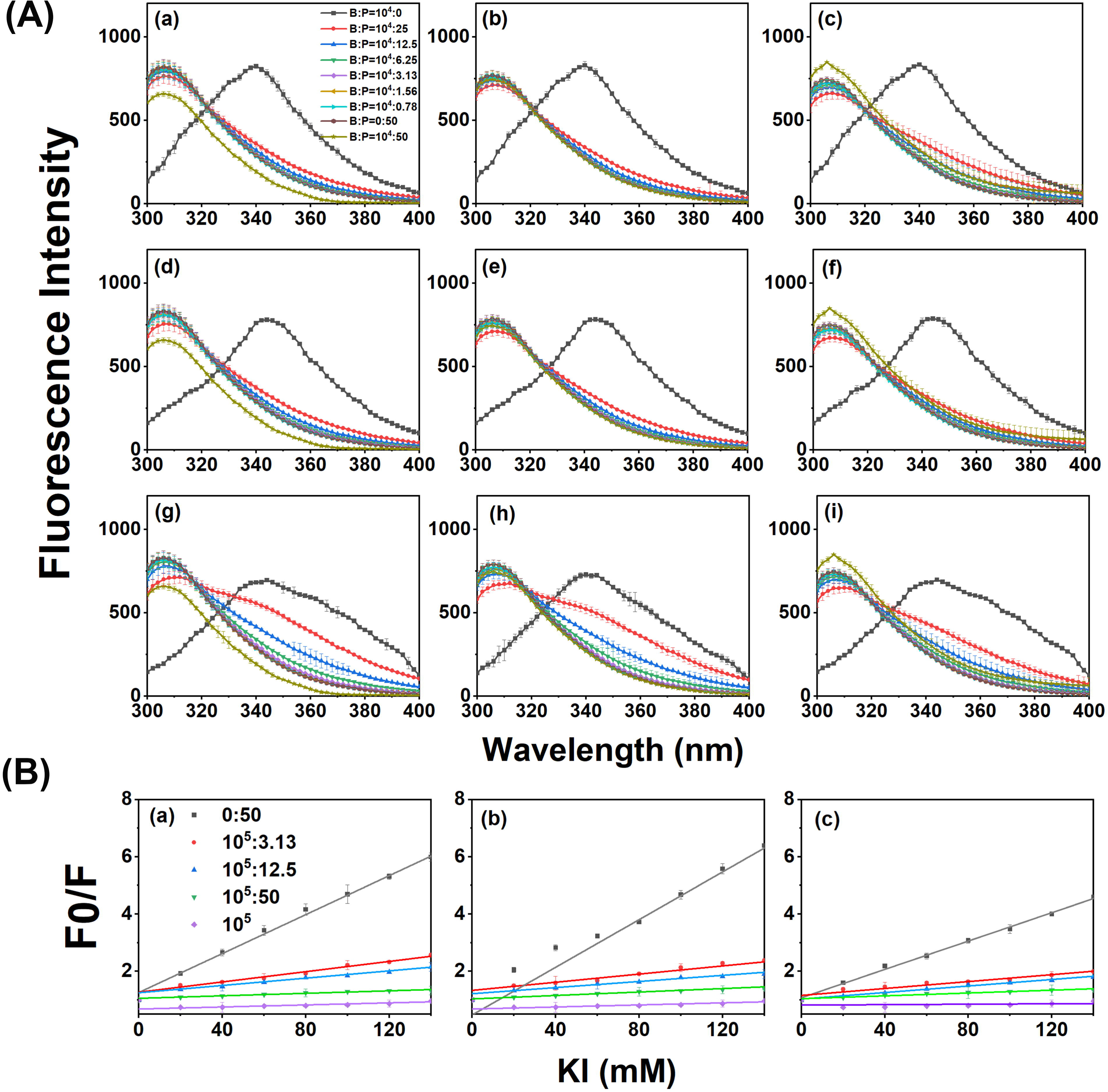
Trp-fluorescence spectra of I4W, L12W, and I1WL5W in the presence of bacteria. (**A**) a-c: I4W incubated with bacterial suspension for 10 min (a), 1 h (b), and 3 h (c), respectively; d-f: L12W incubated with bacterial suspension for 10 min (d), 1 h (e), and 3 h (f), respectively; g-i: I1WL5W incubated with bacterial suspension for 10 min (g), 1 h (h), and 3 h (i), respectively. (**B**) Stern-Volmer plots for the fluorescence quenching of Trp residues in the presence of bacteria a: I4W; b: L12W; c: I1WL5W. Data are presented as mean ± standard error of the mean (SEM), n = 3.

Stern-Volmer quenching analysis employing aqueous KI solutions validated the lipid bilayer penetration capacity of the tryptophan-rich peptides. Hydrophobic insertion of tryptophan moieties into the membrane cores creates quencher resistance through spatial segregation from KI interactions. The attenuation of fluorescence directly correlated with reduced aqueous-phase tryptophan exposure. Quantitative measurements (**Figure 3B**) revealed maximal fluorescence reductions of 72.9%, 78.4% and 83.7% for I1WL5W, I4W and L12W, respectively, relative to the control baselines.

### 3.5 Effects of Trp-containing AMPs on the membrane permeability of MDRKP 1203 cells

Electrostatic perturbation of the cytoplasmic barriers of MDRKP 1203 by tryptophan-rich peptides was evaluated through membrane depolarization monitoring via potentiometric tracking. The assay utilized DiSC3-5, a voltage-sensitive fluorophore that exhibits self-quenching properties upon intracellular penetration. This cationic fluorochrome undergoes fluorescence attenuation upon cytoplasmic internalization due to self-quenching phenomena. Alterations in membrane integrity provoke dye expulsion from compromised cells, generating quantifiable fluorescence restoration [43]. All three peptides provoked immediate fluorescence intensification, which correlated with dosage escalation and demonstrated that the membrane potential collapsed in treated pathogens (**Figure 4A-C**). At 4× MIC, I4W and I1WL5W achieved 80-90% depolarization relative to the Triton X-100 controls, indicating near-complete membrane destabilization.

**Figure 4.**
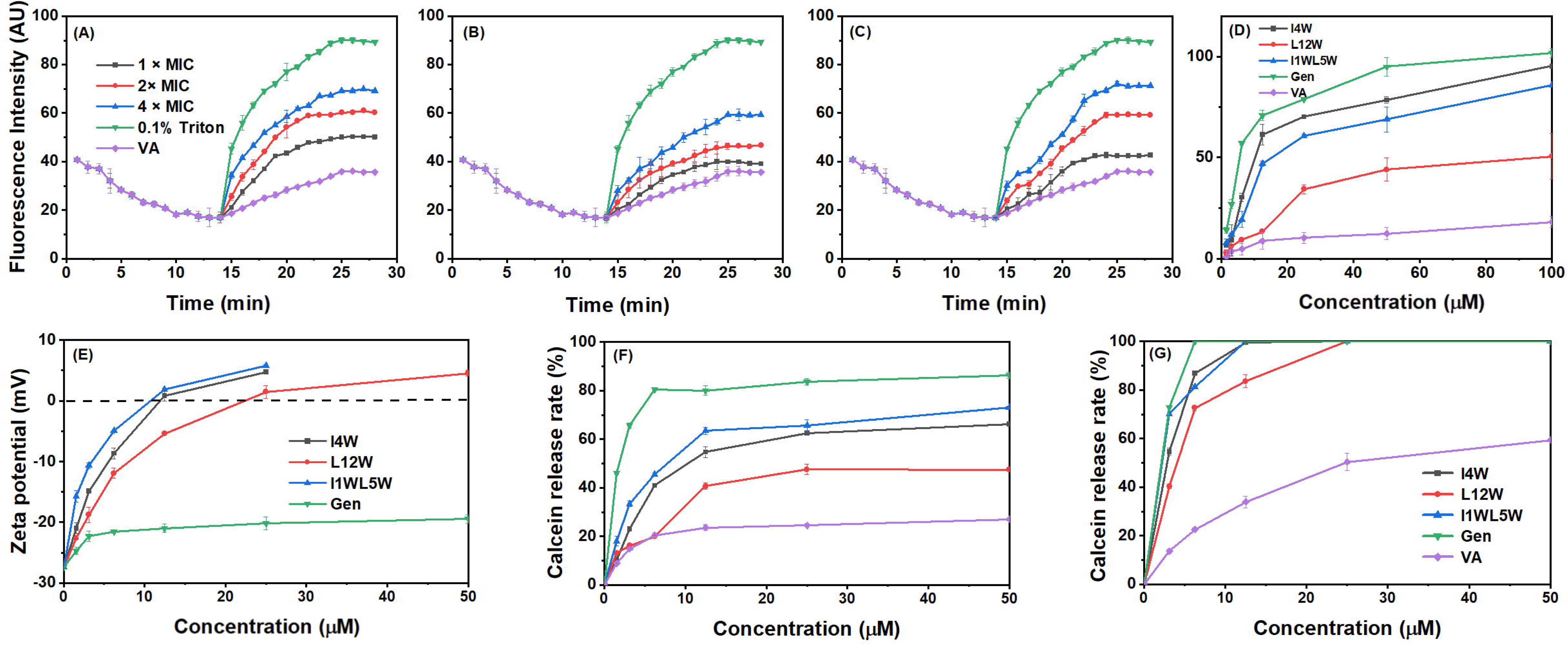
The effect of I4W (**A**), L12W (**B**), I1WL5W (**C**) on MDRKP 1203 bacterial membrane. Peptide concentrations were 1× MIC, n = 3. Effect of three peptides on the outer membrane permeability of MDRKP 1203 cells (**D**), n = 3. Bacterial cells were incubated with 1× MIC of the peptides in the presence of NPN and the fluorescence intensity was detected when the dye was inserted into the outer membrane. Gen was used as the positive control. The effect of I4W, L12W, I1WL5W to the surface charge of MDRKP 1203 cells (**E**), n = 3. The calcein release from the liposomes after the addition of 1× MIC concentrations of three peptides. Liposome compositions: (**F**) (PE/PG/PC: 0.15:1.85:1, w/w/w) and (**G**) PC/PG/PE/LPS (5:2.95:1.05:1, w/w), n = 3. Data were expressed as the mean ± SEM.

The membrane permeabilization dynamics of three tryptophan-enriched peptides against MDRKP 1203 cellular barriers were investigated through NPN fluorochrome interrogation. The hydrophobic 1-N-phenylnapthylamine probe exhibits fluorescence amplification exclusively upon outer membrane destabilization by permeabilizing agents [44]. Quantitative fluorometric analyses demonstrated concentration-gradient-dependent signal enhancement across all the tested peptides (**Figure 4D**). Comparative evaluation revealed that the I1WL5W and I4W peptides induced superior membrane destabilization relative to the L12W peptide, although both exhibited reduced disruptive capacity compared with the benchmark performance of gentamicin.

The zeta potential was measured to monitor the surface charges on the outer membrane of MDRKP1203 cells. Electrostatic alterations in MDRKP 1203 cellular envelopes under peptide exposure were quantified through zeta potential analysis (**Figure 4E**). Incremental administration of tryptophan-containing peptides induced surface charge transition from −27.36 mV at baseline to +5.23 mV at maximal displacement. Membrane charge reversal (>0 mV) occurred at the 12.5 μM threshold for both I1WL5W and I4W, demonstrating complete electrostatic counteraction of native bacterial membrane negativity. The L12W peptide necessitated a doubled concentration (25 μM) to achieve comparable surface charge neutralization efficacy. Notably, exposure at 50 μM gentamicin failed to induce measurable charge reversal, resulting in the maintenance of persistent electronegativity across bacterial surfaces. To evaluate the membrane-disruptive properties of tryptophan-rich peptides, a calcein encapsulation-release methodology was implemented. Vesicular models simulating gram-negative bacterial envelopes were constructed. The outer membrane surrogates employed POPC/POPG/POPE/LPS (5:2.95:1.05:1 mass ratio) multilamellar carriers, whereas the inner membrane analogs utilized POPE/POPG/POPC (0.15:1.85:1 mass proportion) multilamellar carriers. Quantitative assessment of vesicular content release due to peptide-induced membrane perturbation was performed via fluorometric measurement of encapsulated dye efflux across reconstituted bilayers. As depicted in **Figure 4F**, I1WL5W and I4W induced rapid calcein leakage from bacterial outer membrane-mimetic LUVs at relatively low concentrations (0–12.5 μM). Specifically, at 6.25 μM, I1WL5W and I4W elevated the calcein release rate to over 40%, whereas L12W only elicited a 20% release. When the concentration increased to 12.5 μM, I1WL5W triggered a release rate exceeding 60%, I4W induced a rate exceeding 50%, and L12W resulted in a 40% release. Upon exceeding 12.5 μM, the release rate exhibited a steady, gradual upward trend; at the maximum tested concentration (50 μM), I1WL5W, I4W, and L12W mediated calcein leakage from bacterial outer membrane-mimetic LUVs at rates of 72.6%, 65.9%, and 47.0%, respectively. Additionally, I1WL5W and I4W also provoked rapid calcein leakage from bacterial inner membrane-mimetic LUVs within the low-concentration range (0–12.5 μM). At 6.25 μM, I4W drove a release rate >85%, I1WL5W induced a rate >80%, and L12W elicited a rate >70%. At 12.5 μM, I1WL5W and I4W achieved a stable 100% release rate, while L12W required a concentration of 25 μM to reach complete (100%) calcein release (**Figure 4G**). Collectively, these results demonstrate that I1WL5W and I4W exert more potent disruptive effects on both bacterial outer and inner membrane-mimetic LUVs compared to L12W. Moreover, the membrane-disruptive activities of these three peptides are significantly superior to that of vancomycin, while being slightly less potent than that of gentamicin.

### 3.6 Effect of the Trp-Containing Peptides on MDRKP 1203 Biofilms

Biofilm inhibition assays revealed concentration-dependent suppression of MDRKP 1203 biofilm formation by Trp-containing peptides (1/4 to 2× MIC, 24 hour). Quantitative analysis revealed 19%, 26%, and 21% inhibition for I4W, L12W, and I1WL5W at 1/4 MIC, respectively, and these inhibition percentages increased to 33%, 27%, and 28%, respectively, at the 1× MIC (**Figure 5A**). The MBIC50 values were determined to be 12.2 μM (I4W), 23.7 μM (L12W), and 12.8 μM (I1WL5W) (**Table 3**). Trp-containing peptides decreased the dispersal of preformed biofilms (1-day-old biofilm) in a dose-dependent manner. As shown in **Figure 5B**, the peptides at concentrations of 1/2 and 1 × MIC reduced the biofilm biomass by 10-12% and 27-39%, respectively. However, the decline in biofilm density was not significant when the peptide concentration was decreased to 1/4 or 1/8 MIC. Compared with the control, these results indicated that low concentrations of Trp-containing peptides had no effect on the dispersal of preformed biofilms. The MBRC_50_ values of I4W, L12W and I1WL5W against 1-day-old MDRKP 1203 biofilms were 43.7, 47.4 and 44.8 μM, respectively.

**Figure 5.**
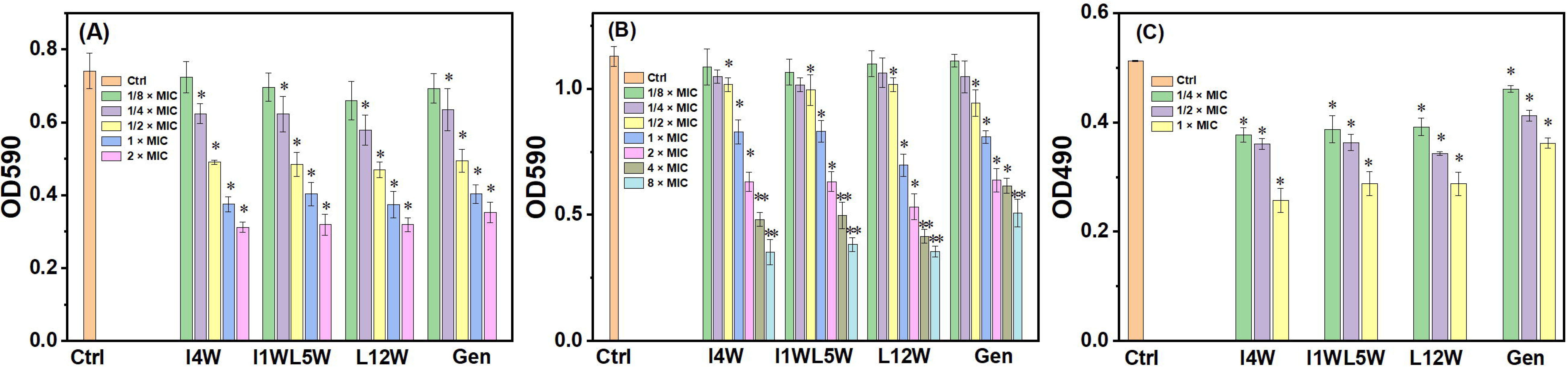
Effect of the Trp-Containing peptides on MRKP 1203 biofilms. (**A**) Inhibition effect of I4W, L12W, I1WL5W on biofilm formation; the MRKP 1203 cells were incubated at 37 °C for 24 h in the absence (control) and presence of a series concentration of the peptides, n = 3. (**B**) Effect of the peptides on 1-day-old biofilm: Biofilms of MRKP 1203 cells were grown for 1 d in the absence of the peptide, and then the 1-day-old biofilm was treated with a series concentration of the peptides at 37 oC for 24 h, n = 3. (**C**) Effect of the Trp-containing peptides on extracellular polysaccharide of MRKP 1203 biofilms. Results represent the means and SD from triplicate experiments, n = 3. Data were expressed as the mean ± SEM by one-way ANOVA; *0.01 < p < 0.05, **p < 0.01, versus control group. *p < 0.05 and **p < 0.01 indicate statistically significant differences between peptide and control.

**Table 3.** MBIC_50_ and MBRC_50_ of peptides against MDRKP 1203 biofilm.

| Peptides | MIC(μM) | MBIC<sub>50</sub> (μM) | MBRC<sub>50</sub> (μM) |
| --- | --- | --- | --- |
| I4W | 12.5 | 12.18 | 43.68 |
| L12W | 25 | 23.66 | 47.37 |
| I1WL5W | 12.5 | 12.84 | 44.77 |
| Gen | 3.13 | 3.26 | 13.72 |

Exopolysaccharides are important components of biofilms that provide protection from antimicrobial agents. We next tested if the Trp-containing peptides at subinhibitory concentrations decrease the formation of biofilms in MDRKP 1203 cells. As shown in **Figure 5C**, I4W, L12W and I1WL5W applied at 1/4 MICs decreased exopolysaccharide production in MDRKP 1203 cells by 20%, 15% and 13%, respectively. When the I4W, L12W and I1WL5W peptide concentration was increased to 1× MIC, exopolysaccharide production was reduced by 60%, 50% and 53%, respectively. The inhibitory effect of gentamicin was inferior to that of the three peptides.

### 3.7 Trp-containing peptides protect mice from MDRKP1203-induced pulmonary infection

An MDRKP 1203-induced lung infection mouse model was constructed to study the therapeutic effect of Trp-containing antimicrobial peptides. The mice received a nasal drip injection of 1 ×10^7^ CFU/mL MDRKP 1203 for one hour, followed by intraperitoneal (i.p.) administration of the I1WL5W Trp-containing peptide at doses of 2, 4, or 8 mg/kg body weight. Saline (control) and gentamicin (2 mg/kg body weight; positive control) were administered as controls. Mice received injections each day over a span of 9 days. Bacterial colonization was measured in the mouse lungs and blood. As shown in **Figure 6A** and **6B**, the bacterial counts significantly increased after one day of bacterial inoculation, reaching 10^4^ CFU/mL in the lung tissues and blood, and I1WL5W reduced the counts in a dose-dependent manner. Mice treated with I1WL5W at a dosage of 8 mg/kg body weight experienced a reduction exceeding 2 log unit CFU over a period of 9 days. In contrast, the control group treated with 2 mg/kg body weight gentamicin presented superior antibacterial effects compared with the other three peptides.

**Figure 6.**
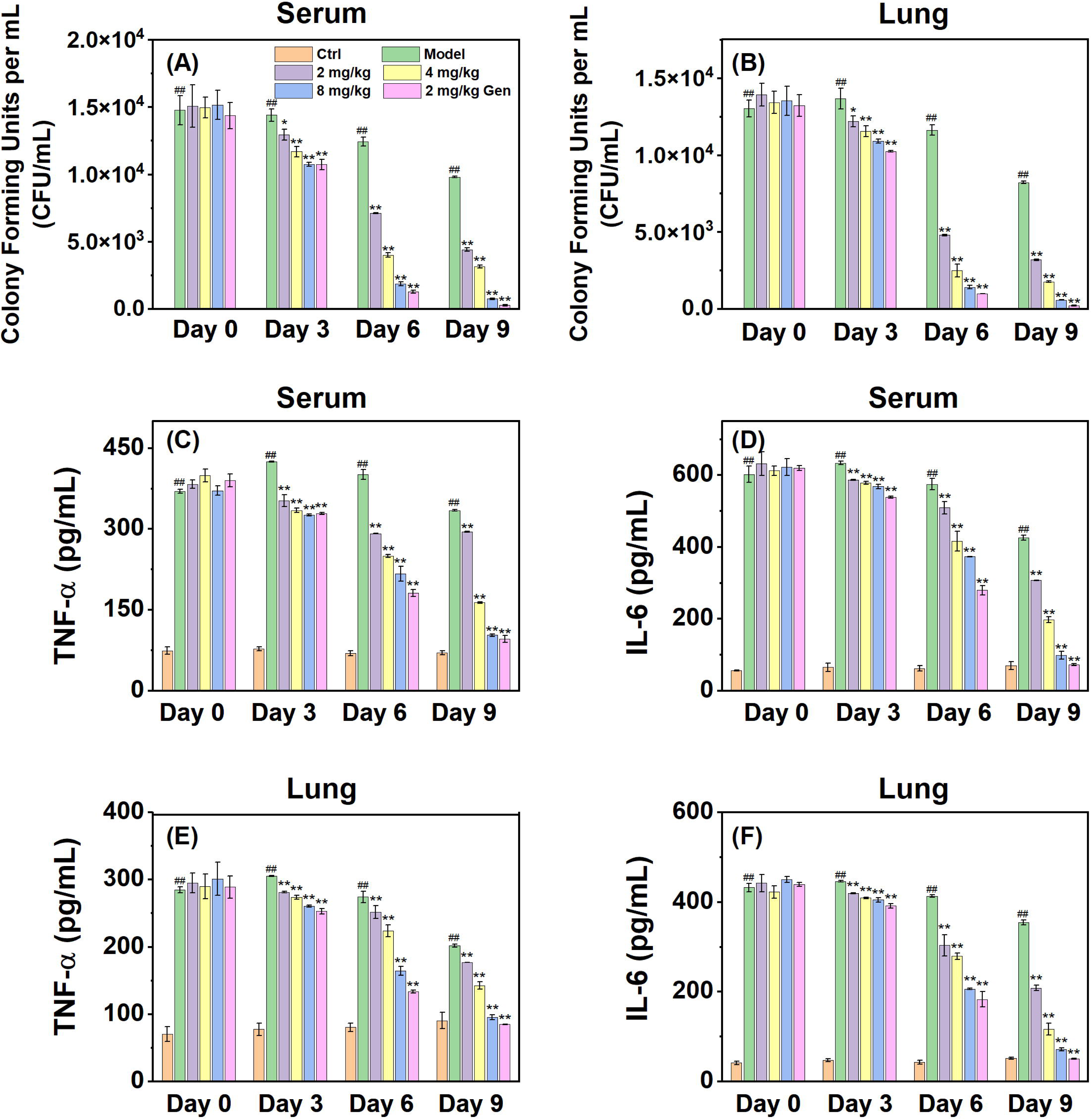
Treatment of peptides against mouse pulmonary infection caused by MDRKP 1203. The number of bacterial CFU in the pulmonary infection of mice treated with MDRKP 1203 using the spiral-plating method, and bacterial colonization were counted in blood (**A**) and lung (**B**) of mice (n = 3); The release of the proinflammatory cytokines TNF-α (**C**, **E**) and IL-6 (**D**, **F**) in mouse lungs and blood was measured at 3, 6 and 9 d (n = 3). Data were expressed as the mean ± SEM by one-way ANOVA; ##*p* < 0.01 versus control group; *0.01 < *p* < 0.05, \*\**p* < 0.01, versus model group.

Systemic and pulmonary inflammatory responses were quantified through TNF-α and IL-6 measurements in serum and lung homogenates. MDRKP 1203-challenged mice presented significant cytokine elevation (p<0.01), which was attenuated by I1WL5W in a dose- and time-dependent manner. As shown in **Figure 6C** and **D**, the serum TNF-α/IL-6 levels decreased by 17.1–23.3%/7.4–10.4% (2–8 mg/kg, Day 3), 31.4–49.0%/19.7–41.2% (Day 6), and 33.5–89.7%/51.6–87.5% (Day 9), respectively. Similarly, the pulmonary TNF-α/IL-6 levels were decreased by 7.9–14.6%/6.1–9.3% (Day 3), 27.5–40.1%/32.0–53.7% (Day 6), and 42.0–83.9%/53.3–86.3% (Day 9) as shown in **Figure 6E** and **F**. These results demonstrated dose-responsive suppression of infection-driven cytokine cascades.

Histopathological analysis revealed marked pulmonary pathology in MDRKP 1203-infected mice, characterized by alveolar interstitial congestion, vascular dilatation, and dense inflammatory infiltrates (**Figure 7**). Compared with no treatment, therapeutic administration of I1WL5W (8 mg/kg) for 6 days attenuated inflammatory pathology, resulting in reduced neutrophilic infiltration and intra-alveolar exudates. After 9 days of treatment, the pulmonary architecture exhibited restored alveolar integrity with transparent parenchyma, minimal residual exudates, and resolution of vascular congestion, indicating near-complete resolution of infection-induced tissue damage.

**Figure 7.**
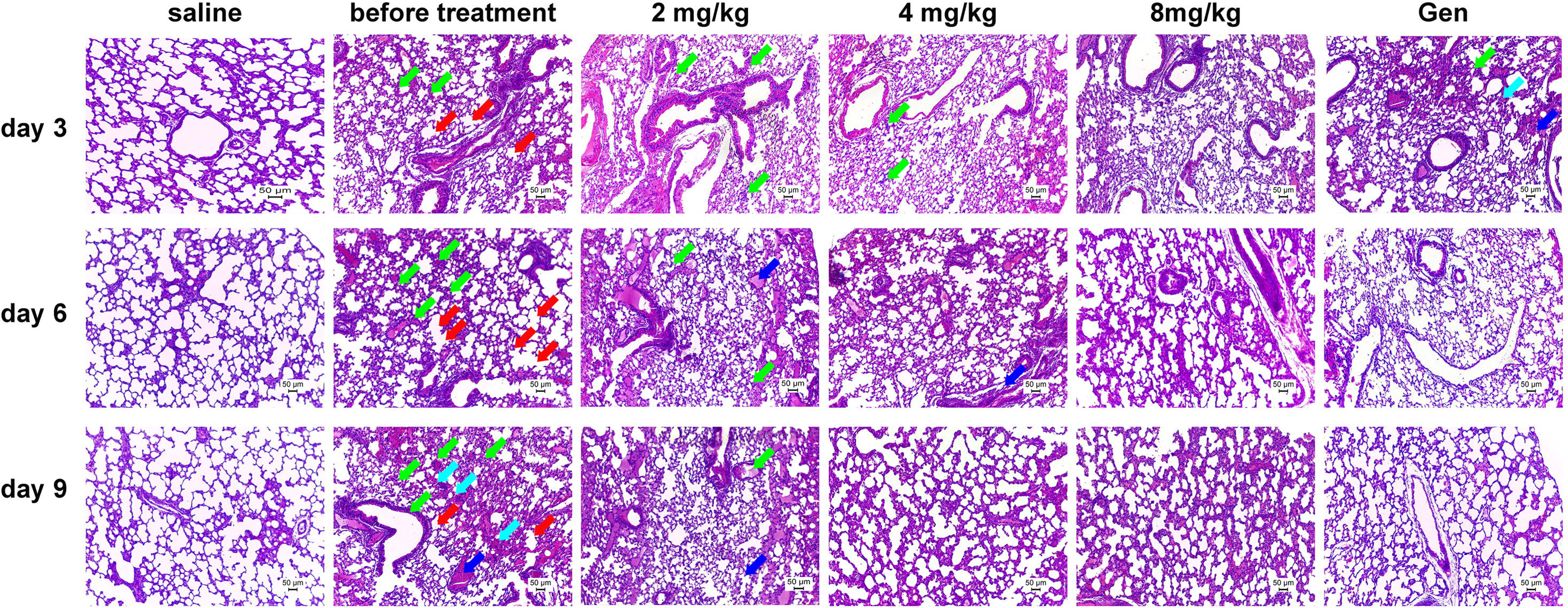
Histological observation of mice lung tissue stained by H&E after treatment of MDRKP 1203. Green arrows indicate inflammatory cell infiltration; Red arrows indicate alveolar wall destruction and enlargement of alveolar spaces; Cyan arrows indicate intra-alveolar hemorrhage; Blue arrows indicate vascular dilation and congestion. Scale bar: 50 μm.

## 4. Discussion

In recent years, owing to the widespread use and overuse of antibiotics, *K. pneumoniae* strains have exhibited increasing resistance to these drugs, rendering their infections challenging to treat[45]. Extensive research has confirmed that clinically isolated *Klebsiella pneumoniae* strains exhibit multidrug resistance to multiple antibiotic classes, specifically aminoglycosides, penicillin derivatives, and carbapenem compounds[46, 47]. This phenomenon is particularly pronounced in hospital settings. Therefore, the development of new antibiotics holds paramount significance. AMPs are recognized as a new category of powerful antimicrobial agents, because of their wide-ranging effects and distinct membrane processes. Additionally, AMPs exhibit significant inhibitory effects on select cancer cells while causing minimal harm to mammalian cells. Importantly, AMPs are less prone to induce resistance than current antibiotics[48, 49].

Prior investigations from our laboratory have established the potent broad-spectrum antimicrobial efficacy of tryptophan-modified antimicrobial peptides (Trp-AMPs) against gram-negative and gram-positive pathogens[21, 22, 38, 41]. The present results revealed that several Trp-containing AMPs had significant bactericidal effects on the MDRKP 1203 strain, with MIC values ranging from 12.5-50 μM. I4W and I1WL5W had more significant bactericidal effects compared with L12W. The potency represents a marked improvement over the parent peptide, temporin-1CEb. which previously showed no antibacterial activity (MIC =175μM) against *susceptible K. pneumoniae* strains[23]. Importantly, the engineered peptides are not only highly effective against the susceptible strain (MIC = 25 μM for I4W and I1WL5W)[21, 41] but also demonstrate equal or superior efficacy (MIC = 12.5-25 μM) against the MDRKP 1203 strain. The dual effectiveness against both susceptible and resistant pathogens underscores their potential as broad-spectrum antimicrobial agents. In addition, the bactericidal kinetics tests revealed that all the AMPs achieved complete microbial eradication within 180 minutes at 16 × MIC. Rapid bacteriolytic kinetics correlated with prolonged postantibiotic effects, manifesting sustained PAE durations of 4 h after exposure for 60 minutes at 1×MIC. The sustained suppression exceeds the conventional PAF duration of 1-2 hours[50–52], which indicates an irreversible antibacterial action.

To elucidate the structure-activity relationship of the Trp-containing peptides, we employed AlphaFold2 for structural prediction. As depicted in Figure S2, the parental peptide Temporin-1CEb was predicted to predominantly adopt a minimal helical conformation, with its N-and C-terminal regions exhibiting a tendency towards disorder. In contrast, the engineered peptides I4W, L12W, and I1WL5W all exhibited a propensity to form α-helical structures. Notably, the most potent peptides, I1WL5W, was predicted to form well-defined, stable amphipathic α-helices. In the I1WL5W model, the two tryptophan residues at positions 1 and 5 are positioned on the same hydrophobic face of the helix, creating a prominent aromatic/hydrophobic patch (Figure S1). This configuration is ideal for insertion into the lipid bilayer of bacterial membranes. The structure of L12W, while helical, suggests that the C-terminal Trp-12 substitution might slightly distort the helical stability or amphipathicity compared to I1WL5W and I4W. These structural predictions strongly support the hypothesis that the enhanced antibacterial efficacy of I1WL5W and I4W is intrinsically linked to their stabilized amphipathic α-helical conformation, which facilitates optimal interaction with and disruption of bacterial membranes.

Antimicrobial peptides (AMPs) exert their bactericidal activity through the following biphasic mechanism: initial electrostatic attraction between cationic peptide domains and anionic phospholipid headgroups in microbial membranes, followed by hydrophobic insertion mediated by aromatic side chain-bearing residues into the acyl chain regions of the lipid bilayer. This sequential interaction cascade facilitates peptide partitioning into the membrane interior, culminating in structural destabilization[53]. Direct evidence of this mechanism was provided by ultrastructural analyses (SEM and confocal microscopy), which confirmed that our Trp-containing AMPs induce cytoplasmic membrane disruption and eventual bacteriolysis. Notably, I1WL5W and I4W demonstrated potent anionic charge neutralization capacities at submicromolar concentrations, inducing near-neutral zeta potential values (≈0 mV) through targeted electrostatic interactions with bacterial membrane surfaces. To directly assess the impact of the AMPs on membrane permeability, depolarization assays were conducted on bacterial plasma membranes[54, 55]. The biophysical assay results demonstrated mechanistic concordance with the zeta potential measurements, revealing that Trp-containing AMPs initiate bactericidal activity through electrostatic adsorption to microbial surfaces, followed by membrane charge neutralization and subsequent transmembrane potential dissipation.

Trp residues are uniquely known for their ability to interact with the membrane interfacial region, thereby facilitating peptide anchoring to the lipid bilayer[19, 56]. The Trp environment of the peptide can reflect the interactions of the peptides with the cell membrane, which can be detected by fluorescence spectroscopy[32, 57]. Specifically, as peptides transition from an aqueous buffer into the hydrophobic interior of the bacterial membrane, their Trp emission maximum shifts to a shorter wavelength—a phenomenon known as a blue shift. The depth of tryptophan residue integration into lipid bilayers can be quantified through characteristic blue shifts in fluorescence emission maxima, indicative of indole group relocation into hydrophobic membrane microdomains with reduced environmental polarity[30]. It is noteworthy that the observed fluorescence signals unequivocally originate from the Trp residues in the administered peptides rather than from intrinsic bacterial membrane proteins. This specificity is attributed to two key factors: first, the high concentration of peptides (0.75–50 μM), each containing one or more strongly fluorescent Trp residues, ensures that the peptide-derived signal dominates over the relatively weak and constant background fluorescence from bacterial proteins; second, the pronounced blue shifts and quenching profiles represent dynamic changes triggered by peptide-membrane insertion, which cannot be accounted for by the static fluorescence of endogenous bacterial components. The present fluorescence results demonstrated that the blue shift values of the three selected AMPs exceeded 26 nm, indicating the complete localization of tryptophan residues within the hydrophobic environment and their full insertion into the bacterial membrane. Notably, I1WL5W exhibited the most significant blue shift, suggesting that its deep burial results in stronger binding affinity toward the bacterial membrane. Tryptophan fluorescence quenching assays employing KI as a collisional quencher demonstrated reduced solvent accessibility of indole moieties in I4W, L12W, and I1WL5W, indicating deep positioning of Trp residues within the hydrophobic core of MDRKP 1203 membranes. For the majority of gram-negative bacteria, the external membrane serves as a significant shield against the lethal impact of antibiotics[58]. Experiments involving the release of calcein, in which AMPs with tryptophan residues aimed at liposomes replicate the inner and outer membranes of MDRKP 1203’s, revealed that these AMPs fully compromised the integrity of the inner cell membrane at reduced concentrations, leading to 100% release of calcein. Nonetheless, the percentage of calcein secreted from the outer membrane of MDRKP 1203 remained below 80%, even at elevated AMP concentrations. These results are consistent with those from the outer membrane permeability test, showing that AMPs with Trp cause alterations in the permeability of the outer membrane of MRKP 1203 cells, compromising their structural integrity, although this impact is somewhat mild. Importantly, this reduction in membrane permeability is correlated with resistance to antimicrobial peptide killing in vitro because it largely prevents the entry of AMPs into MDRKP 1203 cells. Overall, the present findings suggested that tryptophan residues possess potent membrane-disrupting activity, thereby conferring the unique ability of Trp-containing AMPs to interact with the bacterial cell membrane surface and potentially enhance their antimicrobial efficacy. The Trp-containing peptides developed in the present study have potential as innovative agents for antibiotics and in developing groundbreaking treatment approaches.

The formation of biofilms in bacteria represents a defensive mechanism employed by these microorganisms to adapt to their surrounding environment and shield themselves from the bactericidal effects of antibiotics[59]. Biofilms are widely prevalent in *K. pneumoniae* communities and are considered crucial for the robust pathogenicity and clinical drug resistance of this species[60]. Shielded by biofilms, *K. pneumoniae* exhibits slower growth, diminished metabolism, insensitivity to external stimuli, and significantly reduced antibiotic permeability. These attributes facilitate the evasion of antibiotic killing effects, as antibiotics that are effective against planktonic bacteria often are ineffective against the same bacterial strain in the presence of biofilms, making biofilms pivotal factors in bacterial resistance[61]. The present study demonstrated that minimal levels of I4W, L12W, and I1WL5W successfully prevented the development of MDRKP 1203 biofilms, in a concentration-dependent manner. The suppressive impact intensified as the peptide concentration increased. Extracellular polysaccharides (EPSs) are crucial components of bacterial biofilms and play a significant role in their formation. The EPS-based biofilm matrix consists primarily of nucleic acids, proteins, lipids, and exopolysaccharides[62]. Degrading EPSs and eliminating this protective layer can effectively reduce antibiotic resistance in harmful bacteria[63]. In the present study, I4W, L12W, and I1WL5W each inhibited the EPS component of MDRKP 1203 biofilms. Moreover, these inhibitory effects became more pronounced with increasing peptide concentration.

Furthermore, histological analysis of lung tissues revealed that mice infected with MDRKP 1203 exhibited significant pathological damage to the lungs, characterized by interalveolar space congestion, pulmonary blood vessel dilation, infiltration of numerous inflammatory cells, and partial destruction of the alveolar structure. Notably, the structural integrity of mouse lung tissues after treatment resembled that observed in normal mice. The present findings strongly suggest that I1WL5W has the potential to augment bacterial clearance within the pulmonary system while enhancing host defense against respiratory infections in vivo.Lysis of bacterial cells releases lipopolysaccharide (LPS), a structural component of gram-negative outer membranes, which activates the NF-κB signaling cascade via Toll-like receptor 4 (TLR-4) engagement on immune cells, driving elevated expression of proinflammatory mediators[64, 65]. Compared with the saline control treatment, I1WL5W administration significantly attenuated TNF-α and IL-6 concentrations in the blood and pulmonary tissue homogenates. The maximal suppression of cytokine levels occurred after administering 8 mg/kg I1WL5W for 9 days, which achieved the same suppression as that resulting from gentamicin treatment and approached the baseline concentrations observed in the non-infected cohorts. These results suggested that I1WL5W effectively neutralizes the LPS present on MDRKP 1203 cell membranes, thereby down-regulating inflammatory cytokine expression. Unlike genetically homogeneous inbred strains (e.g., C57BL/6 or BALB/c), the inherent genetic diversity of KM mice more accurately parallels human population variability, which is valuable for pharmacological evaluation. Furthermore, KM mice have been successfully utilized in prior research for establishing pneumonia models[66–68], validating their suitability for our study. Future studies will validate key findings in C57BL/6 models to enhance translational relevance and alignment with established platforms. Regarding the exclusive use of male animals, the use of male KM mice aimed to reduce variability in immunoregulatory capacity potentially introduced by hormonal cycles in females, consistent with methodology in prior pneumonia models[69, 70].

However, several limitations of this study should be considered. Firstly, a comprehensive assessment of the long-term in vivo toxicity, potential hemolytic activity at higher therapeutic doses, and detailed pharmacokinetic/pharmacodynamic (PK/PD) profiles of these peptides remains to be fully elucidated. Future work must prioritize the following assessments to determine the therapeutic potential and clinical translatability of these peptides: (1) Safety & Toxicology: Determine the maximum tolerated dose (MTD) and no observed adverse effect level (NOAEL) in repeated-dose studies, and rigorously evaluate hemolytic activity against human erythrocytes at high doses to define a clear safety window. (2) PK/PD Profiling: Characterize key pharmacokinetic parameters (e.g., half-life, clearance, bioavailability) and establish the critical PK/PD index (e.g., AUC/MIC) that correlates with efficacy in vivo. Secondly. by inducing neutropenia via cyclophosphamide, we established an “immune-deficient” environment mimicking high-risk clinical populations, thereby directly detected the anti-bacterial effects of Trp-containing AMPs. This approach is a well-established and widely adopted standard method for creating neutropenic models to study infections caused by drug-resistant Gram-negative bacteria, including *K. pneumoniae*[71, 72]. It reliably replicates the critical immune deficiencies observed in the target patient population receiving immunosuppressive therapies or with underlying immunocompromising conditions. However, evaluating Trp-containing AMPs in immunocompetent animals is essential for comprehensively understanding their therapeutic potential, particularly for broader pneumonia patient populations. Future work will establish the immunocompetent murine models of MDRKP 1203-induced pneumonia to systematically evaluate the therapeutic efficacy of Trp-containing AMPs. This comparative approach across immune contexts will provide critical data for determining optimal clinical applications.

Bacterial cell membranes represent crucial targets for the development of novel antimicrobial agents against multidrug-resistant microorganisms, as pathogens undergo gradual changes in membrane composition over time[30]. Trp-containing antimicrobial peptides (I4W, L12W, and I1WL5W) exhibited pronounced membrane translocation capacity in multidrug-resistant *K. pneumoniae* (MDRKP 1203), demonstrating high-affinity interactions with lipopolysaccharide (LPS) moieties within gram-negative outer membrane constituents facilitating drug entry into the cells. Therefore, Trp-containing AMPs, particularly I1WL5W, hold promise as potential lead compounds for developing innovative antibiotics that target bacterial membranes to combat multidrug-resistant *K. pneumoniae*.

## Conclusion

Exploring how Trp-containing AMPs function in multidrug-resistant *K. pneumoniae* MDRKP 1203, their impact on biofilm formation and growth, and their therapeutic effectiveness in vivo is vital for identifying prospective antimicrobial drug targets. The present study indicated that AMPs containing Trp residues, especially I1WL5W with two Trp residues, affect mainly the internal and external membranes of bacteria, showing notable antibiofilm capabilities. Furthermore, the present findings demonstrate that I1WL5W can effectively treat MDRKP 1203-induced pulmonary infections in mice. The therapeutic mechanism involves bacterial clearance and growth inhibition, leading to the inhibition of inflammatory responses. This research offers fresh insights into the development of innovative and potent anti-infective AMP molecules aimed at combating bacterial aggressiveness.

## Author contributions

**Fengquan Jiang**: Data curation, Writing–Original draft. **Yanjun Ma**: Data curation, Investigation, Methodology. **Yunfei Zhang**: Methodology. **Dejing Shang**: Supervision, Validation, Writing - review & editing. **Weibing Dong**: Resources, Conceptualization, Funding acquisition, Writing-Reviewing and Editing.

## Acknowledgements

This work was supported by grants from the National Natural Science Foundation of China (grant numbers 32270513 and 32070440), Dalian Science and Technology Innovation Fund (grant number 2021RJ04), the Scientific Research Fund of Liaoning Provincial Education Department (grant number LJ212410165064), and Liaoning Normal University (grant number 25GDL009).

