## Supplemental Data for "Tryptophan-substitution antimicrobial peptide temporin-1CEb: in vitro and in vivo antibacterial activity against clinically isolated multidrug-resistant *Klebsiella pneumoniae*"

**Multidrug-resistant of the other bacterial strain in the emergency ICU (eICU)**

Among the 57 *Staphylococcus aureus* strains examined, a high prevalence of resistance was observed to erythromycin, clindamycin, and penicillin, with penicillin resistance exceeding 80%. Notably, these strains remained susceptible to furantoin, quinupristin/dalfopristin, rifampicin, linezolid, vancomycin, tigecycline, and daptomycin. Moreover, 39 strains of Enterococcus faecalis presented high rates of resistance to erythromycin, ciprofloxacin, rifampicin, penicillin, tetracycline and levofloxacin. S. aureus and S. epidermidis are highly resistant to commonly used antibiotics, with rates of resistance ranging from 56.5 to 100% to oxacillin, erythromycin, ciprofloxacin, clindamycin, penicillin, and levofloxacin (Table S3).

**Table S1** Distribution of cultures and types of pathogenic microorganisms in emergency ICU.

| Type of culture | Culture-positive |
| --- | --- |
|  | ICU [N(%)] |
| blood culture | 156（7.59%） |
| sputum culture | 1555（75.71%） |
| Bronchial lavage fluid culture | 140（6.82%） |
| urine culture | 105（5.11%） |
| secretion culture | 51（2.48%） |
| Other Cultures | 47（2.29%） |
| Total | 2054（100%） |
| Type of microorganism |  |
| Gram-positive bacteria | 182（8.86%） |
| Gram-negative bacteria | 1335（65.00%） |
| Fungus | 537（26.14%） |
| Total | 2054（100%） |

**Table S2** Distribution of pathogenic microorganisms in emergency ICU[N(%)]

| Items | ICU(n=2054) | Total |
| --- | --- | --- |
| **Gram-positive bacteria** |  | 182(8.86%) |
| *S. aureus* | 2.78% |  |
| *E. faecium* | 1.90% |  |
| *E. faecalis* | 0.05% |  |
| *S. hominis* | 1.36% |  |
| *S. epidermidis* | 1.12% |  |
| *S. haemolyticus* | 0.29% |  |
| Others | 1.36% |  |
| **Gram-negative bacteria** |  | 1335(65%) |
| *A. baumannii* | 19.43% |  |
| *E. coli* | 3.51% |  |
| *B. cepacia* | 2.04% |  |
| *P. mirabilis* | 1.51% |  |
| *K. pnenmoniae* | 20.59% |  |
| *S. maltophilia* | 6.86% |  |
| *E. cloacae* | 1.12% |  |
| *C. indologenes* | 0.15% |  |
| *A. junii* | 0.05% |  |
| *H. influenzae* | 0.05% |  |
| *P. aeruginosa* | 6.23% |  |
| *E. aerogenes* | 0.68% |  |
| *S. marcescens* | 0.63% |  |
| *K. oxytoca* | 0.39% |  |
| Others | 1.75% |  |
| **Fungus** |  | 537(26.14%) |
| *C. albicans* | 12.66% |  |
| *C. tropicalis* | 3.26% |  |
| *C. Parapsilosis* | 2.00% |  |
| *C. glabrata* | 7.40% |  |
| *C. krusei* | 0.58% |  |
| Others | 0.24% |  |

**Table S3** Resistance rate of important pathogenic microorganisms in emergency ICU to commonly antibiotics (%)

| Name of antibiotics | **Gram-positive bacteria** | | | |
| --- | --- | --- | --- | --- |
|  | *S. aureus*  (57 strains) | *E. faecium*  (39 strains) | *S. hominis*  (28 strains) | *S. epidermidis*  (23 strains) |
| Benzoxacillin | 31.5 | ND | 89.29 | 91.3 |
| Furantoin | 0 | 4.75 | 5.56 | 0 |
| Sulfamethoxazole | 12.2 | ND | 50 | 60.8 |
| Erythromycin | 73.6 | 94.8 | 92.8 | 65.2 |
| Ciprofloxacin | 45.6 | 100.00 | 76.1 | 73.9 |
| Clindamycin | 73.6 | ND | 85.71 | 56.5 |
| Quinupristin/Dalfopristin | 0 | 0 | 2.3 | 0 |
| Rifampicin | 0 | 76 | 7.14 | 13 |
| Linezolid | 0 | 0 | 0 | 0 |
| Moxifloxacin | 43.8 | ND | 64.29 | 47.8 |
| Penicillin | 82.4 | 100.00 | 100.00 | 100.00 |
| Gentamicin | 7.02 | ND | 17.86 | 21.7 |
| Tetracyclin | 21 | 66.6 | 25 | 30.4 |
| Tigecycline | 0 | 0 | 0 | 0 |
| Vancomycin | 0 | 5.1 | 0 | 0 |
| Levofloxacin | 45.6 | 100.00 | 67.8 | 82.6 |
| Daptomycin | 0 | 0 | 0 | 0 |
|  | **Gram-negative bacteria** | | | |
|  | *E. coli*  (72 strains) | *A. baumannii*  (399 strains) | *K. pnenmoniae*  (423 strains) | *P. aeruginosa*  (128 strains) |
| Amikacin | 5.5 | 57.8 | 11.3 | 3.9 |
| Ampicillin | 92.86 | ND | ND | ND |
| Aztreonam | 61.1 | ND | 77.5 | 35.1 |
| Polymyxin | 0 | 0.25 | 0 | 0 |
| Cotrimoxazole | 59.7 | 69.1 | 44.9 | ND |
| Ciprofloxacin | 70.8 | 96.2 | 70.6 | 48.4 |
| Piperacillin/Tazobactam | 30.5 | 96.59 | 73.2 | 24.77 |
| Gentamicin | 36.1 | ND | 40.11 | 60 |
| Tigecycline | 0 | 1.28 | 0.97 | ND |
| Cefoperazone/Sulbactam | 25 | 38.54 | 71.36 | 16.4 |
| Ceftriaxone | 69.4 | ND | 77.27 | ND |
| Ceftazidime | 52.7 | 94.9 | 73.5 | 21.8 |
| Cefepime | 69.4 | 91.7 | 79.6 | 7.03 |
| Cefuroxime | 70.8 | ND | 78.07 | ND |
| Cefotetan | 14.06 | ND | 26.38 | ND |
| Ampicillin/Sulbactam | 76.56 | ND | 78.03 | ND |
| Tobramycin | 20.8 | 82.2 | 34.7 | 28.9 |
| Imipenem | 16.6 | 95.57 | 72.5 | 40.6 |
| Levofloxacin | 62.5 | 88.9 | 54.1 | 57.8 |
| MeropeneM | 9.38 | 95.79 | 68.61 | 35.29 |
|  | **Fungi** | | | |
|  | *C. albicans*  (260 strains) | *CandidaNDglabrata*  (152 strains) | *C. tropicalis*  (67 strains) | *C. Parapsilosis*  (41 strains) |
| Amphotericin B | 0 | 0 | 0 | 0 |
| 5-Fluorocytosine | 0 | 0 | 0 | 0 |
| Voriconazole | 0 | 22.37 | 17.91 | 12.1 |
| Itraconazole | 0 | 7.89 | 0 | 4.8 |
| Fluconazole | 0 | 7.2 | 19.4 | 24.3 |

ND: Data was not determined

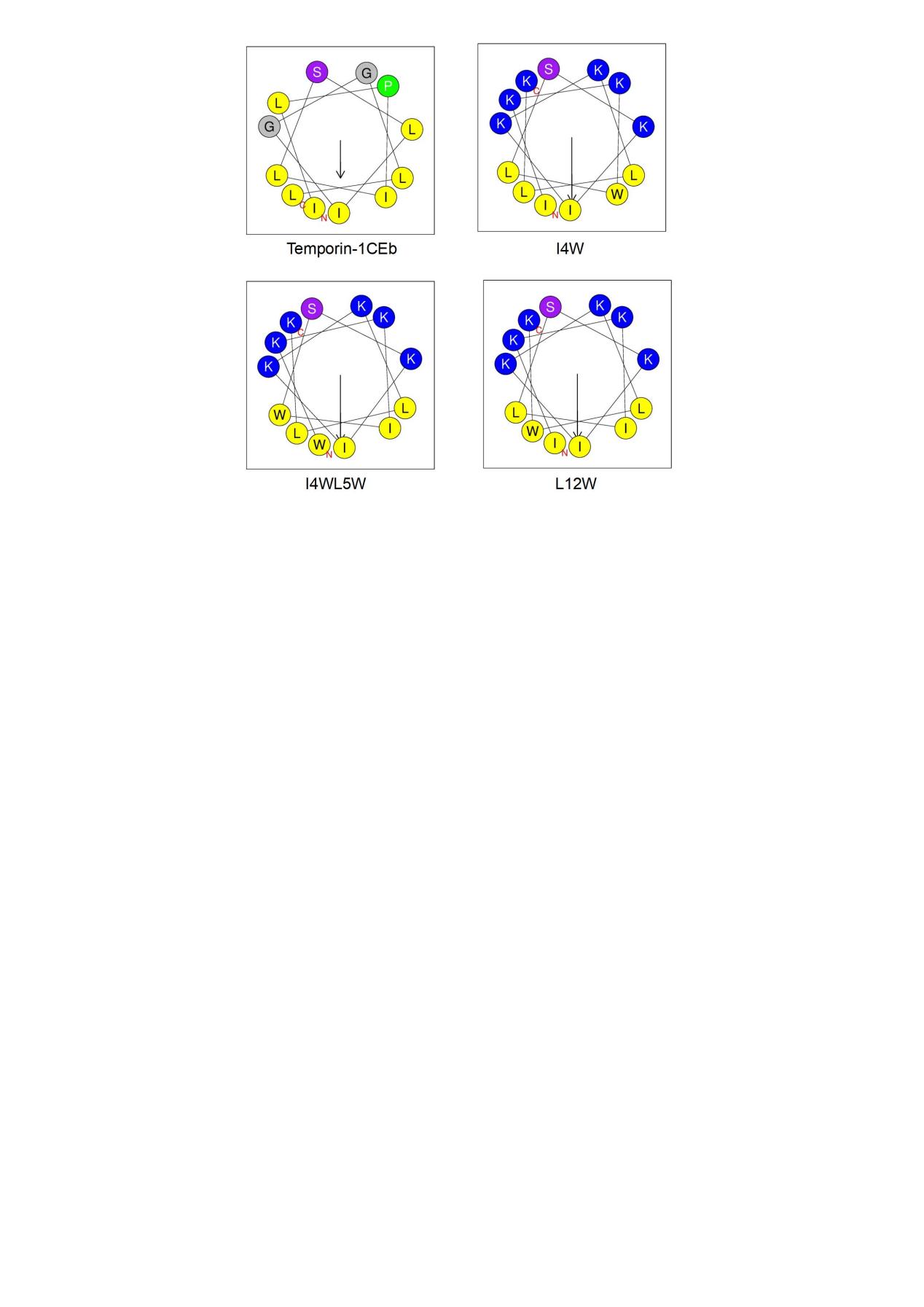

**Figure S1** Helical wheel projections of the parent and engineered peptides. Projections for Temporin-1CEb and the modified peptides (I4W, I4WL5W, and L12W) were generated using HeliQuest. The diagrams are colored according to standard residue properties: blue for basic, red for acidic, purple for polar uncharged, yellow for hydrophobic, and green for special residues (e.g., Gly, Pro).

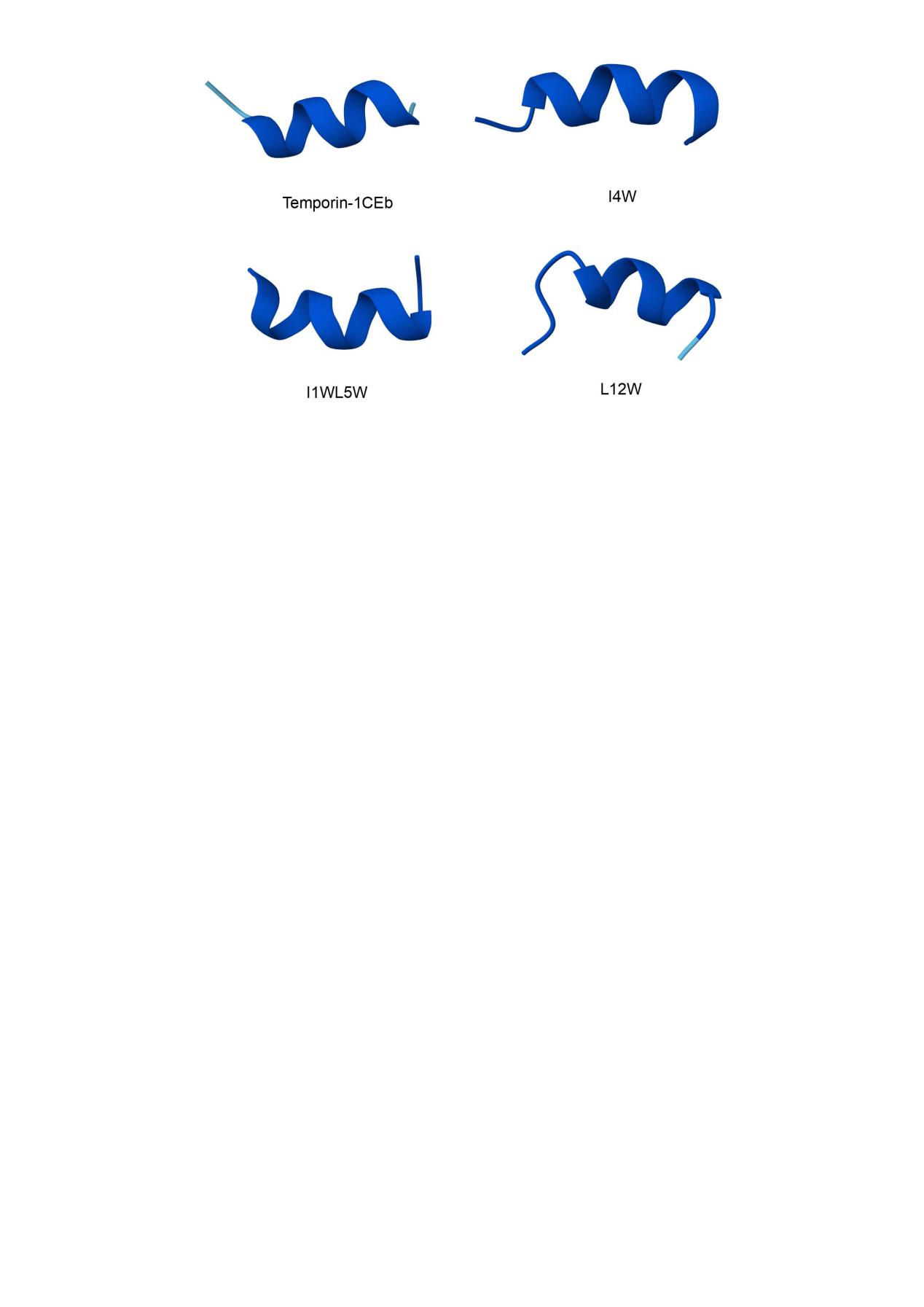

**Figure S2** Predicted structures of the engineered peptides. Three-dimensional models of (A) Temporin-1CEb, (B) I4W, (C) L12W, and (D) I1WL5W were generated by AlphaFold2. Each peptide is depicted as a cartoon representation, colored by the per-residue confidence score (pLDDT). Key tryptophan residues are shown as stick models. The corresponding predicted aligned error (PAE) plots are displayed below each structure.
